# Massive postglacial gene flow between European white oaks uncovered genes underlying species barriers

**DOI:** 10.1101/246637

**Authors:** Thibault Leroy, Quentin Rougemont, Jean-Luc Dupouey, Catherine Bodénès, Céline Lalanne, Caroline Belser, Karine Labadie, Grégoire Le Provost, Jean-Marc Aury, Antoine Kremer, Christophe Plomion

**Affiliations:** BIOGECO, INRA, Univ. Bordeaux, 33610 Cestas, France; Département de biologie, Institut de Biologie Intégrative et des Systèmes (IBIS), Université Laval, GlV 0A6, Québec, Canada; INRA Université de Lorraine UMR 1137 ‘Ecologie et Ecophysiologie Forestières’, route d’Amance, 54280 Champenoux, France; CEA · Institut de Biologie François Jacob, Genoscope, 2 rue Gaston Crémieux, 91057 Evry, France

**Author notes:** **Corresponding author:** Antoine Kremer, INRA, UMR1202 BIOGECO, F-33610 Cestas, France.

**Keywords:** Genome scan, approximate Bayesian computation, demographic inferences, intrinsic and ecological barriers, reproductive isolation

## Abstract

Oaks are dominant forest tree species widely distributed across the Northern Hemisphere, where they constitute natural resources of economic, ecological, social and historical value. Hybridization and adaptive introgression have long been thought to be major drivers of their ecological success. Thus, the maintenance of species barriers remains a key question, given the extent of interspecific gene flow. In this study, we scanned the genomes of four European white oak species for reproductive barriers. We identified the ecological and phylogenic relationships of these species and inferred a long-term strict isolation followed by a recent and extensive postglacial contact. Then, we made use of the tremendous genetic variation among these species (31 million SNPs) to identify genomic regions for reproductive isolation. A literature-based functional annotation of the underlying genes highlighted important functions driving the reproductive isolation between these sister species. These functions were consistent with their ecological preferences and included tolerance to biotic and abiotic constraints. This study holds important implications for the renewal of European forests under global warming.

## Main

Oaks are a diverse group of about 350 to 500 species widely distributed throughout the Northern Hemisphere [1, 2]. The variability in the number of recorded oak species highlights the challenge of delineating species limits within a genus displaying a high degree of morphological diversity, sometimes described as a “botanic horror” by taxonomists [3–5]. Genetic markers have corroborated these taxonomic concerns, particularly in European white oaks, which have been the subject of a large number of genetic surveys. Studies based on nuclear DNA markers have reported unambiguously high levels of admixture between European white oak species, confirming the reported taxonomic issues for oaks [6]. Several detailed empirical studies based on chloroplast. DNA markers have revealed an absence of private chlorotypes between European white oak species, but congruent associations between chlorotypes and expansion routes during the last postglacial recolonization, suggesting cytoplasmic capture via recurrent hybridization and backcrossing [7, 8]. Recent advances in oak genomics [9, 10] have made it possible to investigate interspecific gene flow at the whole-genome scale. Indeed, Leroy et. al. [11] have provided evidence suggesting that extensive secondary contacts have occurred between four European white oak species, probably at start of the current interglacial period. These results reconcile earlier findings of contrasting species differentiation at the nuclear and organelle levels. The inferences drawn are also consistent with the persistence of genomic regions impermeable to gene flow due to functional reproductive barriers, corresponding to a typical case of semi-isolated species [11]. However, the genetic basis of these barriers remains unknown.

Controlled pollination trials have provided empirical evidence for the existence of strong reproductive barriers in these four European white oak species [12, 13]. Ecological preferences *in situ* have also been previously reported, with tolerance to dry (*Q. petraea*) or wet (*Q. robur*) sites (14), or acidic (*Q. pyrenaica*) or limy (*Q. pubescens*) soils [15] but fine-grained ecological surveys do not yet exists for all these four species. The four species occupy different geographic ranges (Fig 1 A): extending up to Scandinavia for *Q. petraea* and *Q. robur*, whereas the other two species are present mostly in Mediterranean and sub-Mediterranean regions. However, the distribution ranges of these species overlap in some areas, mostly in South-West. France, but the four species are rarely found together in the same stand [but see ref. 6]. The overlapping species ranges in South-West. France thus provide an ideal “natural laboratory” [16] for investigating reproductive barriers between these European white oak species.

**Figure 1:**
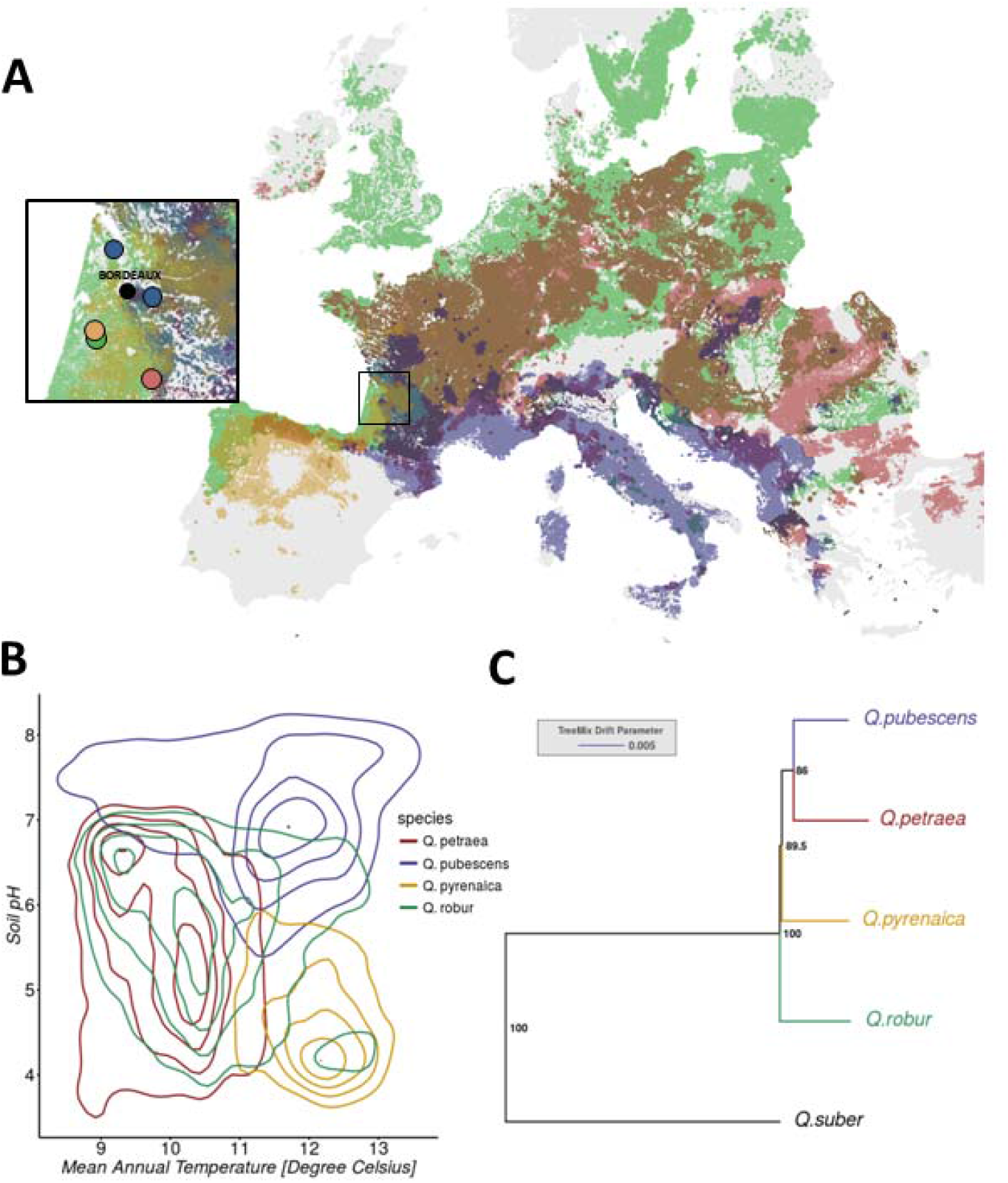
Continental-scale species distributions and origin of the study material (A) and, ecological (B) and phylogenic relationships of the four European white oak species under investigation (C).

Here we combined state-of-the-art methods in population genomics to explore the genomic distribution of reproductive barriers (Fig. 1): (i) we used approximate Bayesian computation (ABC) to perform ascertainment bias-free demographic inferences in order to refine estimates of the timing of secondary contacts, and (ii) scan genomes for reproductive barriers. Our findings identified important intrinsic and ecological functions driving the reproductive isolation of these four oak species including tolerance to biotic and abiotic constraints, and intrinsic mating barriers.

## Results

### Ecological preferences of the four species

We intersected the distribution maps of the four species (Fig. 1A) with climatic and soil data derived from a large-scale floristic survey in France. Bivariate density distributions (Fig. 1B) show clear patterns of ecological preferences among the four white oak species. As expected, the two so called temperate white oaks (*Q. petraea* and *Q. robur*) are more frequently observed under cooler climates than Mediterranean and sub-Mediterranean species (*Q. pubescens* and *Q. pyrenaica*). pH of the soils segregates particularly *Q. pyrenaica* from *Q. pubescens*. Although we could not combine climatic and soil data over the whole species’ ranges, univariate density distributions for both climate (Fig S1) and soil pH (Fig S2) based on continental-scale data showed similar trends.

### Divergence and post-glacial secondary contact between European white oaks

A total of 31,894,340 SNPs were identified after the filtering of variants with a low minor allele frequency (MAF<0.02) in population samples for the four species (*Q. petraea, Q. robur, Q. pubescens, Q. pyrenaica*), corresponding to one SNP every 23.2 bp, on average. We also used genome-wide data for a *Q. suber* accession described by Leroy *et al*. [11] to root a phylogenetic tree and investigate relationships between species for 9,084,835 of the 31.9 million SNPs. The best maximum-likelihood tree suggested that *Q. robur* initially diverged from the ancestor of the other three species (Fig. 1C).

We then randomly selected 50,088 SNPs from the entire set of 31.9 million SNPs for ascertainment bias-free demographic ABC inference. We compared two models of divergence with gene flow (Fig. 2) for each of the six possible species pairs: an isolation-with-migration model assuming constant gene flow since the divergence time (Tsplit), and a model assuming secondary contact with gene flow starting at Tsc, a time point after divergence (Tsc < Tsplit). For all pairs, we obtained strong statistical support for the secondary contact model (>98% posterior probability, Tables 1 & S1), consistent with our previous findings based on individual data for 3,524 SNPs [11].

**Figure 2:**
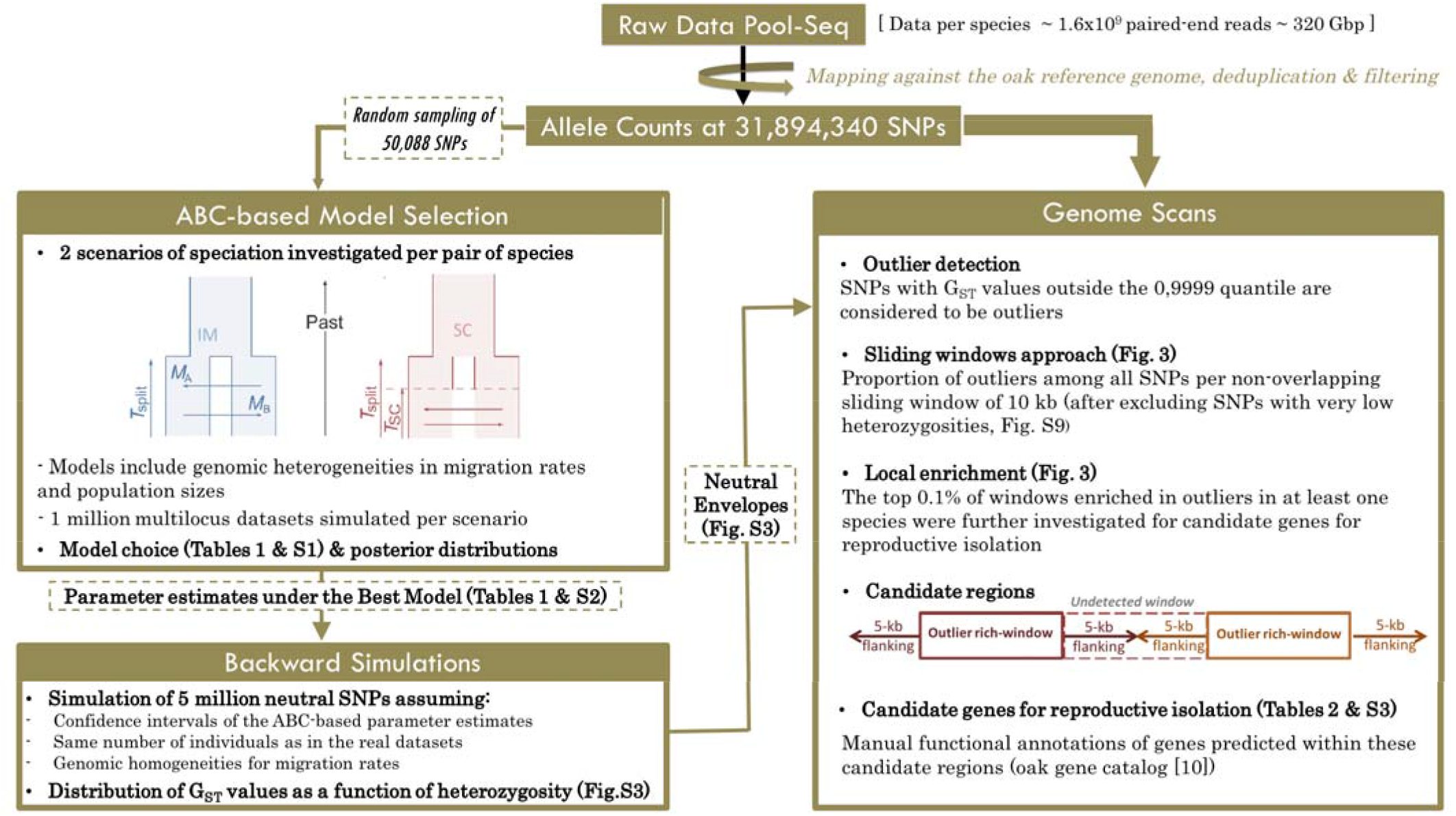
Workflow used to identify genes contributing to reproductive isolation between four European white oak species. A subset of 50 thousand of the called SNPs was selected at random and used for model selection under an ABC framework and the generation of parameter estimates under the best model. Large neutral datasets were then simulated to create null envelopes for the iden tification of SNPs displaying significant departure from expectations under neutrality. We searched for candidate genes in regions enriched in outliers.

**Table 1:** Posterior probabilities of the SC scenario and timing of secondary contacts. Mean (bold) relative posterior probability of the secondary contact scenario and standard deviation (round brackets). Median (bold) and 95% confìdence intervals (square brackets) for both the inferred ratio between divergence time (Tsplit) and time of the secondary contact (Tsc) and the secondary contact ‘expressed in number of years) after setting Tsplit to 10 million years (the upper bound for the divergence of this species complex, [17]). More details are given in Tables S1 & S2.

| Pair | Post. Probability SC | $T_{SC}/T_{SPLIT}$ estimates | $T_{SC}$ (years ago) |
| --- | --- | --- | --- |
| *Q. robur* – *Q. petraea* | **0.98883**<br>(±0.01089) | **1.487 x 10<sup>-3</sup>**<br>[0.43-4.05] x 10<sup>-3</sup> | **14,870**<br>[4,300-40,500] |
| *Q. robur* – *Q. pyrenaica* | **0.99249**<br>(±0.01152) | **1.197 x 10<sup>-3</sup>**<br>[0.31-4.95} x 10<sup>-3</sup> | **11,970**<br>[3,100-40,500] |
| *Q. robur* – *Q. pubescens* | **0.98719**<br>(±0.01342) | **0.245 x 10<sup>-3</sup>**<br>[0.09-0.72} x 10<sup>-3</sup> | **2,450**<br>[900-7,200] |
| *Q. pubescens* – *Q. petraea* | **0.98549**<br>(±0.02231) | **2.176 x 10<sup>-3</sup>**<br>[0.77-6.24] x 10<sup>-3</sup> | **21,760**<br>[7,700-62,400] |
| *Q. pubescens* – *Q. pyrenaica* | **0.99087**<br>(±0.01152) | **0.865 x 10<sup>-3</sup>**<br>[0.32-2.16] x 10<sup>-3</sup> | **8,650**<br>[3,200-21,600] |
| *Q. pyrenaica* – *Q. petraea* | **0.99378**<br>(±0.00697) | **0.383 x 10<sup>-3</sup>**<br>[0.10-1.14] x 10<sup>-3</sup> | **3,830**<br>[1,000-11,400] |

We generated parameter estimates under the best-fitting secondary contact model for each pair of species. Taking into account the 95% confidence intervals for each Tsc/Tsplit ratio (Tables 1 & S2) and the divergence time between these species (1-10 million years, [2,17], the analysis yielded quite large estimates with secondary contact occurring between 100 and 62,400 years ago, corresponding to up to 1,225 generations, assuming a generation time of 50 years [18]. Even assuming the upper bound for the divergence between these species (10mya [17]), median estimates of the timing of secondary contacts had much less variation and ranged between 2,450 and 21,760 years (up to 435 generations). These estimates are consistent with the general hypothesis of a resumption of secondary gene flow at the start of the current interglacial period.

### Narrow regions of non-neutral evolution

We then took advantage of these demographic inferences to perform differentiation outlier tests. We performed extensive backward simulations (5,000,000 independent SNPs) under the best inferred scenario to generate null distributions for each pairwise comparison (Fig. S3, see also *Online Results*). The most outlier-enriched windows were retained for the identification of candidate s genes underlying species barriers. We identified 281 windows containing the highest proportion of outliers (top 0.1% of window enriched in outliers for at least one pair of species). We then analyzed the clustering of these outlier-enriched windows. We defined a candidate genomic region by merging close windows, *e.g*. two contiguous sequences of two outlier-enriched windows, with possible interruption by a single undetected window (Fig. 2). The 281 windows were distributed over 215 candidate genomic regions, distributed over all chromosomes (blue lines, Fig. 3).

**Fig. 3:**
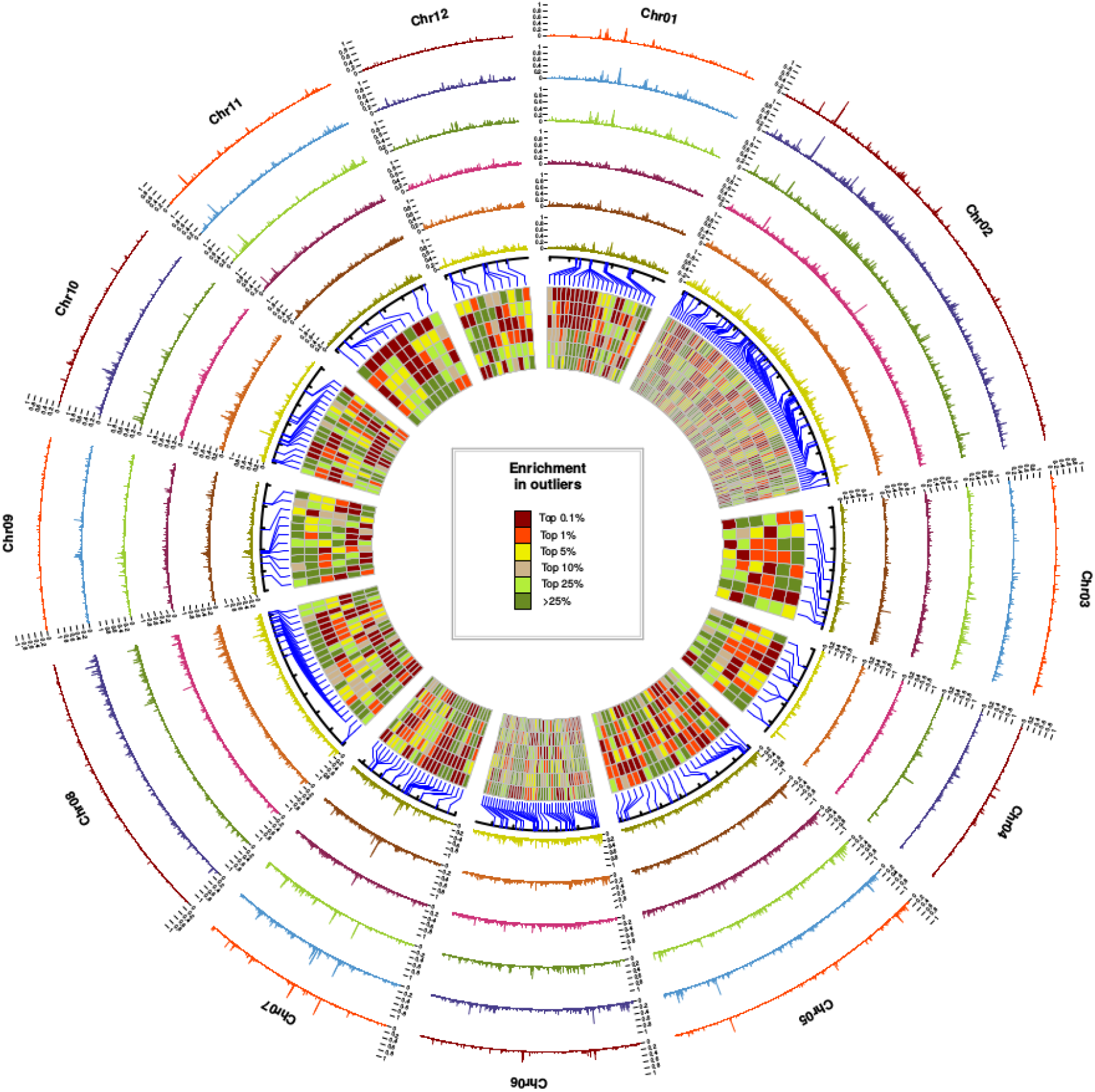
Local density in outliers per non-overlapping 10 kb sliding window. From outside to inside, the species pairs are Q. robur/Q. petraea, Q. pyrenaica/Q. petraea, Q. pubescens/Q. petraea, Q. robur/Q. pubescens, Q. robur/Q. pyrenaica and Q. pubescens/Q. pyrenaica. Detailed patterns are accessible from: httpsV/github.com/ThibaultLeroyFr/GenomeScansByABC/ Each rectangle in the inner circle represents the level of enrichment in outliers with He ≤ 0.2 for each pair of species at a given position in the genome, assuming the same order of pairs. These rectangles correspond to the 281 most outlier-enriched windows found in at least one of the six pairs (top 0.1%).

We listed all the *Quercus* genes located within or flanking these 215 genomic regions. We identified 227 genes distributed over 133 of the 215 regions, with very few candidates per region (mean: 1.71±1.76 genes per region, 1.49 ±0.82 genes after excluding 5 regions with chloroplast.-like DNA signatures). We subdivided genes into three major functional categories, and particularly focused on 32 candidate genes based on quality and sharpness of the annotations in terms of physiological function (Table 2, see also Sup Info). The first category comprises genes underlying the ecological preferences of the four species: tolerance of water deficit, cold tolerance, adaptation to alkaline soils. The second includes genes involved in biotic interactions, such as immune responses, resistance to biotic stresses, and mycorrhization. The third gatherss to genes probably contributing to intrinsic barriers, and includes genes with functions related to flowering time, pollen recognition, pollen growth and embryo development.

**Table 2:** Genomic positions, gene names, annotations for 32 candidate genes. All other annotations are available in Table S3 (see FileS 1 for details, including references). Species patterns were determined from the analysis of the most outlier-enriched windows between pairs, as detailed in Table S4.

| *chr.* | *positions (regions)* | *Candidate gene* | *Gene name* | *Gene functions* | *#Paralogs* | *Pattern* |
| --- | --- | --- | --- | --- | --- | --- |
| ***Intrinsic barriers*** |  |  |  |  |  |  |
| <u>*Flowering*</u> |  |  |  |  |  |  |
| Chr08 | 62493250-62513250 | Qrob_P0412860.2 | Two-component response regulator-like PRR73 | photoperiodic flowering response, circadian clock | 1 | Complex |
| Chr08 | 62673250-62693250 | Qrob_P0749650.2 | Floral homeotic protein APETALA 2 \| Transcr. factor RAP2-7 | Delay transition to flowering time, biotic/abiotic stresses | 0 | Complex |
| Chr07 | 44592072-44632072 | Qrob_P0088630.2 | Transcr. activator DEMETER (DME) / protein ROS1-like | Transcriptional activator required for floral development | 1 | Complex |
| <u>*Pollen development (Cycloartenol synthases)*</u> |  |  |  |  |  |  |
| Chr06 | 28660694-28680694 | Qrob_P0684330.2 | Cycloartenol synthase 2 | Sterol or triterpenoid synthesis, pollen development | 13 | Complex |
| Chr06 | 28690694-28710694 | Qrob_P0684400.2 | Cycloartenol synthase 2 | Sterol or triterpenoid synthesis, pollen development | 0 | Shared |
| Chr06 | 28770694-28810694 | Qrob_P0684360.2 | Cycloartenol synthase 2 | Sterol or triterpenoid synthesis, pollen development | 0 | Shared |
| <u>*Pollen recognition & seed germination (G-type lectin S-receptor-like Serine/threonine kinase genes)*</u> |  |  |  |  |  |  |
| Chr03 | 26118565-26138565 | Qrob_P0538230.2 | G-type lectin S-receptor-like STK At2g19130 | Putatively involved in recognition of pollen | 0 | Complex |
| Chr05 | 4001785-4021785 | Qrob_P0641650.2 | G-type lectin S-receptor-like STK At1g11300 | Putatively involved in recognition of pollen | 190 | Complex |
| Chr06 | 1453464-1473464 | Qrob_P0430150.2 | G-type lectin S-receptor-like STK LECRK | Regulates expression of immunity genes & seed germination | 36 | **Q. pyrenaica-specific** |
| Chr06 | 1513464-1563464 | Qrob_P0430190.2 | G-type lectin S-receptor-like STK LECRK | Regulates expression of immunity genes & seed germination | 36 | **Q. pyrenaica-specific** |
| Chr08 | 1749735-1769735 | Qrob_P0138480.2 | G-type lectin S-receptor-like STK LECRK | Regulates expression of immunity genes & seed germination | 36 | **Q. pyrenaica-specific** |
| <u>*Embryo development & organogenesis*</u> |  |  |  |  |  |  |
| Chr12 | 30514806-30534806 | Qrob_P0436820.2 | Transcriptional corepressor LEUNIG-like isoform | Leaf, flower (gynoecium) and embryo development | 5 | **All except. Q. pub/Q.pyr** |
| Chr02 | 31611918-31651918 | Qrob_P0297580.2 | CHD3-type chromatin-remodeling factor PICKLE | Repressor of LEC1, activator of embryo development | 0 | Complex |
| Chr06 | 43006752-43026752 | Qrob_P0309300.2 | Probable N-acetyltransferase HLS1-like | Auxin-responsive gene expression, shoot organogenesis | 1 | **All except. Q. rob/Q.pet** |
| Chr08 | 49805801-49825801 | Qrob_P0248780.2 | Receptor-like protein 12 (RLP12) | Meristem maintenance control, organogenesis | 0 | Complex |
| <u>*Photoreceptor & UV-B tolerance*</u> |  |  |  |  |  |  |
| Chr02 | 37788250-37808250 | Qrob_P0338870.2 | Ultraviolet-B receptor UVR8 | Photoreceptor/response to UV, circadian clock, stomata | 2 | Shared |
| Chr03 | 34667252-34687252 | Qrob_P0500290.2 | DNA mismatch repair protein MSH2 | UV-B-induced DNA damage response pathway | 0 | Shared |
| Chr06 | 11400887-11420887 | Qrob_P0577750.2 | Transcription factor MYB12 | Positive regulator of flavonoid biosynthesis, UV-B tolerance | 0 | **All except. Q. rob/Q.pet** |
| ***Ecological barriers - abiotic stresses*** |  |  |  |  |  |  |
| <u>*Nramp metal transporters*</u> |  |  |  |  |  |  |
| Chr09 | 7739510-7759510 | Qrob_P0191830.2 | Metal transporter Nramp5 | Manganese and cadmium uptake | 8 | **Q. pubescens-specific** |
| Chr06 | 38764748-38784748 | Qrob_P0097150.2 | Metal transporter Nramp6 | Involved in iron ion homeostasis | 1 | **All except. Q. rob/Q.pet** |
| <u>*Dehydration/lateral root growth*</u> |  |  |  |  |  |  |
| Chr08 | 68418235-68443235 | Qrob_P0457680.2 | Protein DEHYDRATION-INDUCED 19-like | Stress-induced sensor, interacting with CPK11 & S-Rnase | 0 | **All except. Q. rob/Q.pet** |
| Chr02 | 27061274-27081274 | Qrob_P0304800.2 | Transcription factor WER | Controls cell fate specification, e.g. hairy roots or stomata | 0 | **All except. Q. rob/Q.pet** |
| Chr02 | 28240136-28260136 | Qrob_P0299670.2 | Root Primordium Defective 1 (RPD1) | Lateral root morphogenesis; active cell proliferation | 0 | Complex |
| Chr02 | 36392234-36442234 | Qrob_P0422470.2 | Alkaline/neutral invertase CINV2 | Regulator of root growth, sucrose catabolism | 7 | Shared |
| Chr09 | 19506093-19526093 | Qrob_P0418880.2 | 1-aminocyclopropane-1-carboxylate synthase | Ethylene biosynthetic process, lateral root formation | 4 | **All except. Q. rob/Q.pet** |
| Chr04 | 19182244-19222244 | Qrob_P0652510.2 | Phosphatidylinositol-3-phosphatase myotubularin-1 (or 2) | Role in soil-water-deficit stress | 2 | Shared |
| Chr02 | 96008743-96028743 | Qrob_P0387640.2 | Protein WVD2-like 6 | Organ stockiness (periph. root cap, trichomes & leafs) | 0 | **All except. Q. rob/Q.pet** |
| <u>*Freezing/cold adaptation*</u> |  |  |  |  |  |  |
| Chr09 | 18536093-18556093 | Qrob_P0768740.2 | Dehydration-responsive element-binding protein 1 | Key role in freezing tolerance and cold acclimation | 6 | Complex |
| Chr02 | 112153196-112173196 | Qrob_P0339800.2 | B3 domain-containing transcription factor VRN1 | Vernalization responsiveness, repressor of FLC | 3 | **All except. Q. pub/Q.pyr** |
| ***Ecological barriers - biotic stresses*** |  |  |  |  |  |  |
| <u>*Pathogen resistance/mycorrhization*</u> |  |  |  |  |  |  |
| Chr10 | 12310682-12340682 | Qrob_P0070130.2 | Transportin MOS14 | Plant immunity (splicing of resistant genes) | 0 | **Q. pyrenaica-specific** |
| Chr01 | 26763631-26783631 | Qrob_P0648170.2 | Ubiquitin carboxyl-terminal hydrolase 12-like | Protein deubiquitination, regulator of disease resistance | 3 | Complex |
| Chr05 | 3053435-3073435 | Qrob_P0622110.2 | Nodulation-signaling pathway 1 (NSP1) | Nodulation & mycorrhization | 0 | Complex |

### Species-specific ecological and non-ecological reproductive barriers

Unlike studies aiming at interpreting every region enriched in outliers, our objective was rather to focus on genes displaying distinct patterns among pairs of species. This is especially important since these situations are unexpected to arise via background selection (see *Online Results* for details). After excluding genes with a “shared” pattern, several different situations were observed (Table S4): (i) 9 regions enriched in outliers for all but one pair of species (including 7 regions for all pairs except *Q. robur – Q. petraea* and 2 regions for all pairs except *Q. pubescens – Q. pyrenaica* pairs), (ii) 5 regions specific to all pairs sharing the same species (4 for *Q. pyrenaica* and 1 for *Q. pubescens)* and (iii) 11 regions with more complex patterns.

Among the nine regions with an “all·versus-one-pair” relationship, seven excluded the *Q. robui/Q. petraea* pair and the other two excluded the *Q. pubescens / Q. pyrenaica* pair. Four of the seven genes for which the *Q. robur/ Q. petraea* pair was excluded are known to be involved in drought tolerance or in lateral root growth (Table 2). This pattern is consistent with the higher drought tolerance of *Q. pyrenaica* and *Q. pubescens* compared to *Q. petraea* or *Q. robur* [19], Reciprocally, we observed an “all vs. one” pattern (undetected for the *Q. robur/Q. petraea* pair) for a VRN1 gene involved in responsiveness to vernalization and known to play a key role in cold acclimation in many plant species [20]. Overall, the genomic variation of these nine genes parallels the Northern-Southern distribution of the studied species, suggesting that the underlying barriers are driven by climate preferences (Fig 1 A, B).

Among the five genes with branch-specific patterns, *i.e*. found in all pairs containing a given species, four have *Q. pyrenaica*-specific patterns and concerned genes involved in plant immunity, including three encoding G-type lectin S-receptor-like serine/threonine kinases (LECRKs) and one encoding a transportin (MOS14). *Q. pyrenaica* is extremely sensitive to oak powdery mildew, a pathogen that was introduced into Europe at the start of the 20th century [21]. Soon after the first detection of the fungus in Europe, high mortality rates were reported for *Q. pyrenaica* in the humid warm-temperate forests of Southwestern and Western France [21 and references therein], *Q. pyrenaica*-specific alleles at several genes involved in immunity may, therefore, be the signature of the high susceptibility of *Q. pyrenaica* to biotic stresses in moist environments. Additionally, we identified a *Q. μubescens*-specific pattern for a gene encoding a metal transporter (Nrampδ) involved in the assimilation of manganese and cadmium in rice and barley [22]. Manganese assimilation is known to be essential for many plant functions, but manganese availability in the soil tends to decrease with increasing pH, and becomes limiting beyond a soil pH of 6.5. This gene probably signs the greater ecological preference of *Q. pubescens* for lime-rich soils (Fig 1B) in comparison to the other three species.

Among genes with complex patterns, we identified many candidate genes for intrinsic premating and postmating barriers. We identified several genes involved in the timing of flowering, including APETALA2 and PRR73. APETALA2 is a key transcription factor for the establishment of the floral meristem [23]. Similarly, PRR73 contributes to flowering time variation in barley and wheat [24 and references therein]. Several species-specific genes may also be involved in mating barriers, including previously described ecological genes with pleiotropic effects, such as *VRN1*. In addition to its primary role in vernalization, *VRN1* is involved in the repression of *FLC* (itself a known repressor of flowering) in *Arabidopsis*, through a vernalization-independent, floral pathway [20]. Several of the candidate genes for intrinsic barriers identified are involved in pollen or embryo development, suggesting that both premating and postmating intrinsic barriers operate in oaks. Three of these genes encode cycloartenol synthases known to be essential for pollen development in *Arabidopsis* [25].

## Discussion

The increasing availability of genomic resources for phylogenetically related species has the potential to greatly improve our understanding of their evolutionary trajectories and the molecular basis of their reproductive isolation as shown here for European temperate oaks. Our demographic reconstruction supports long periods of isolation between these oak species for most of their history leading to the gradual loss of shared alleles and the accumulation of reproductive barriers. We further found evidence of a systematic shift in their trajectories that occurred towards the beginning of the Holocene. More precisely, this shift took place while the oak species were migrating northwards as the climate became warmer, and resulting in their encountering in central Europe. In line with our previous conclusions [11]. our inferences cannot however exclude that, a few secondary contact periods had already taken place earlier. Still, the mixture of different species and populations in central Europe occurring during the Holocene was so massive that private (or near private alleles) were redistributed among interfertile species. Indeed, current levels of interspecific differentiation are extremely low along almost all the genome (mean interchromosomal 10kb estimates of Fst below 0.08 for all pairs), at a level compatible with many reports of within-species population structure in the literature [26]. However, at some narrow regions distributed throughout the genome, interspecific differentiation reaches extremely high levels (10-kb estimates of Fst above 0.8). These peaks most likely correspond to narrow regions where selection counteracted the homogenizing effect of gene flow, thus leading to the present-day highly heterogeneous landscape of differentiation.

The highest differentiated SNPs contributing to reproductive barriers mostly set apart Southern (*Q. pyrenaica* & *Q. pubescens*) from Northern species (*Q. robur* & *Q. petraea*). While this observation is inconsistent with the inferred phylogeny (ref. 11; Fig. 1C), it however coincides with the climatic preferences of the four species (Fig 1 A,B). We also found genetic support for *Q. pubescens* preference for alkaline soils. When comparing the genomic footprints of differentiation between *Q. pyrenaica* and the other three species, we found genes involved in plant immunity, in line with previous reports for higher mortality rates in this species due to pathogens [21]. Finally our results also show that, peaks of differentiation never reach fixation, even in regions exhibiting the strongest reproductive barriers. The possibility for a very low permeability to gene flow at these barriers therefore calls for more research about the maintenance and evolution of reproductive isolation between European white oaks.

Overall, our results suggest that key selective abiotic and biotic factors triggered by post glacial environmental changes have molded the extant landscape of species reproductive barriers in European temperate oak species. We anticipate that these drivers will operate during ongoing climate changes as Mediterranean oak species (*Q. pyrenaica* & *Q. pubescens*) are migrating northwards getting in contact in more Northern latitudes with local temperate species (*Q. petraea* and *Q. robur*).

## Methods

### Ecological niche of the four species

#### French data

We delineated the extant ecological niche of the four oak species in France (Fig S4) by using their distribution maps based on the National Forest Inventory and climatic data extracted from the Chelsa data base [27]. In addition to the climatic data we added pH values of the soil. Proxies of pH values were derived from floristic data the National Forest inventory floristic plots installed since 2005. Floristic composition of these inventory plots was compared to existing database to calculate proxys of pH values [28]. We intersected the distribution maps with the climatic rasters (30” resolution) and calculated a 2D density plot of species presence in the climatic (mean temperature and precipitation) and pH domain using the R package “ggplot.2” v. 2.2.1 [29] under R v. 3.2.2

#### European data

European distributions maps were constructed based on presence data of species made available by the European atlas of forest tree species [30]. Climatic data are based on the Chelsa database [27] and soil pH were derived from JCR data [31]. Since both data origin from different geographical sites, we only computed univariate density distributions using ggplot2, after using a similar procedure than for the French data.

### Sampling and sequencing

We sampled populations of the four *Quercus* species in stands of natural origin located in South-West. France. We sampled 13 *Q. petraea* trees in Laveyron (Landes, France), and 20 *Q. robur* and 20 *Q. pyrenaica* trees from the Landes EVOLTREE “Intensive Study Site” (ISS). We also sampled 18 *Q. pubescens* trees from two sites in Gironde: 12 in Branne and 6 in Blaignan (Gironde, France) (see Table S5 for details). *Online Methods* contains detailed information on the methodologies used from DNA extraction to sequencing. Overall, between 1,617,465,418 and 1,813,403,677 reads per pool were retained for analysis, corresponding to 313 Gb (425X) to 356 Gb (483X) of raw data. Raw data have been deposited in the Sequence Read Archive (SRA): ERP105626.

### Mapping and calling

All reads were then mapped against the v2.3 oak haplome assembly [10]. with bowtie2 v. 2.1.0 [31]. using standard parameters for the “sensitive end-to-end” mode. PCR duplicates were removed with Picard v. 1.106 (http7/broadinstitute.github.io/picard/). Samtools v.1.1 [32] and Popoolation2 v. 1.201 [33] were then used to call biallelic SNPs with at least 10 alternate alleles and a depth between 50 and 2000X at each position. To ensure a reasonably low rate of false positives due to Illumina sequencing errors, all SNPs with a MAF lower than 0.02 were discarded. We obtained allele counts for a total of 31,894,340 SNPs. Sliding-window Fst were calculated from allele frequencies with the popoolation2 bioinformatics software suite [33].

### Demographic inferences & genome scans

We used a strategy combining ABC [11, 26] and backward simulations to scan genomes for reproductive barriers (Fig. 2). The ABC approach explicitly takes into account confounding effects of barriers to gene flow [34] and linked selection [26, 35] on demographic inferences. This was done by modeling among loci variation in (i) effective migration rate to take into account barriers to migration [36] and (ii) effective population size to take into account linked selection [37]. *Online Methods* contains detailed information on the methodologies used for demographic inferences and genome scans.

### Functional annotations

For all genes found within regions enriched in outliers, we conducted BlastP searches in both the SwissProt and nr protein databases. Only BlastP results with e-values lower than 10e-5 were considered for protein function annotation. After identification of the protein function by BlastP analyses, extensive manual literature searches were performed. We also reported information from a previous identification of orthologous and paralogous genes in 16 plant species, including *Q. robur*, performed with OrthoMCL (see ref. 10, for details).

## Acknowledgments

This research was funded by the French ANR (GENOAK project, 11-BSV6-009-021) and by the European Research Council under the European Union’s Seventh Framework Programme (TREEPEACE project, FP/2014-2019; ERC Grand Agreement no. 339728). We thank the Genotoul Bioinformatics Platform Toulouse Midi-Pyrenees (Bioinfo Genotoul) and the Biogenouest BiRD core facility (Université de Nantes) for providing computing and storage resources. We also thank Jorge A. P. Paiva for providing access to *Q. suber* data and Camille Roux for fruitful discussions concerning ABC. We would like to thank fellow members of the pedunculate oak genome consortium for helpful advice and suggestions.

